## Supplementary_Information for "Kalium channelrhodopsins effectively inhibit neurons in the small model animals"

### Supplementary methods

#### *C. elegans* constructs

gBlocks (IDT) containing codon optimized cDNAs encoding either one of the opsins (ACR1, KCR1-ET, KCR1(GS) or KCR2-ET) that was tagged with YFP at its C-terminus (with three synthetic introns to enhance expression) was PCR amplified and ligated in the KpnI and EcoRI sites of *sdf-9P::mCherry* vector using the following primer sets:

ACR1-f and ACR-1-YFP-r for *sdf-9p::ACR1::YFP*; KCR-1-f and KCR-1-YFP-r for *sdf-9p::KCR1::YFP*; KCR-1-GS-f and KCR-1-GS-YFP-r for *sdf-9p::KCR1(GS)::YFP*; KCR-1-f and KCR-1-YFP-r for *sdf-9p::KCR2::YFP*.

Genomic DNA corresponding to pan-neuronal promoter (*snt-1p*) was PCR amplified from *C. elegans* genome using the following primers, snt-1P-FseI-F and snt-1P-AscI-R, and then ligated in the Fsel and Ascl sites, to generate *snt-1p::ACR1::YFP*, *snt-1p::KCR1::YFP, snt-1p::KCR1(GS)::YFP,* and *snt-1p::KCR2::YFP*. The plasmid was co-injected with *elt-2::mCherry* at 10ng/µl each into the gonads of adult N2 hermaphrodites by microinjector (InjectMan 4) to establish transgenic strains.

#### Live worm imaging by confocal fluorescence microscopy

For imaging experiments, L4 hermaphrodite worms were transferred to a glass slide and immobilized on 3% agarose pads using 2-3µl 1mg/µl levamisole diluted in M9 buffer. Images were then captured under a 100x objective. Multiple transgenic lines of each transgene were examined for fluorescent expression and localization patterns. Spinning disc confocal (SDC) microscopy was performed on a setup built around a Nikon Ti2 inverted microscope equipped with a Yokogawa CSU-W1 confocal spinning head, a Plan-Apo objective (100 × 1.45 NA), and a back-illuminated sCMOS camera (Prime 95B; Photometrics). Excitation light was provided by 488 nm/150 mW (Coherent) (for YFP) (power measured at optical fiber end) through DPSS laser combiner (iLAS system; Gataca systems). All image acquisition and processing were controlled by MetaMorph (Molecular Device) software. Images were acquired with exposure times in the 400–500 ms range.

**Supplementary table 1. Summary of genotypic and statistical information for all experiments.**

The Figure and Panel columns index the respective information to the figures in the manuscript. The Genotype 1 and Genotype 2 columns provide the genotype for the respective *Test* and *Control* genotypes. The Assay and Protocol columns contain information on the assay used and the type of experiment performed. The sample size (N) columns display the total number samples used in each experiment and the experimental iterations (N iterations) columns display the number of repetitions for the respective experiment. The estimation statistics columns show the mean difference effect size delta (Δ Effect size) between the control and test genotypes with 95% confidence intervals (CI) with the corresponding *P* value (P).

**Table S2: *C. elegans* genotypes and DNA constructs**

| **Strain genotype** | **ID** |
|---|---|
| *C. elegans: yasEX257 [snt-1p::ACR1::YFP (10ng/ul); elt-2P::mCherry (10ng/ul)]* | SAH743 |
| *C. elegans: yasEX259 [snt-1p::KCR1::YFP (10ng/ul); elt-2P::mCherry (10ng/ul)]* | SAH745 |
| *C. elegans: yasEX260 [snt-1p::KCR2::YFP (10ng/ul); elt-2P::mCherry (10ng/ul)]* | SAH746 |
| *C. elegans: yasEX261 [snt-1p::KCR1(GS)::YFP (10ng/ul); elt-2P::mCherry (10ng/ul)]* | SAH747 |

| **Recombinant DNA constructs** | **ID** |
|---|---|
| *snt-1p::ACR1::YFP* | RAI77 |
| *snt-1p::KCR1::YFP* | RAI78 |
| *snt-1p::KCR1(GS)::YFP* | RAI79 |
| *snt-1p::KCR2::YFP* | RAI80 |
| *sdf-9p::mCherry* | JB164 |

| **Recombinant DNA sequences** | **ID** |
|---|---|

| | |
|---|---|
| acccttgGCTAGCgtcgacGGTACCggtagaaaaaATGAGTTCCATCACGTGCGACCCTG<br>CCATCTACGGCGAGTGGTCCCGAGAGAACCAGTTCTGCGTAGAGAAGTCCTTGAT<br>AACCCTGGATGGAATTAAGTACGTCCAGCTGGTGATGGCAGTCGTGTCAGCATGT<br>CAAGTCTTTgtaagtttaaacagttcggtactaactaaccatacatatttaaattttcagTTCATGGT<br>GACAAGAGCACCGAAGGTTCCTTGGGAAGCGATTTATTTGCCGACCACCGAAAT<br>GATTACCTATTCATTGGCCTTTACGGGAAATGGTTACATTCGAGTCGCTAATGGCA<br>AGTATCTTCCGTGGGCTCGAATGGCATCTTGGCTTTGCACCTGCCCTATAATGCTT<br>GGACTTGTATCCAACATGGCCTTAGTAAAGTACAAATCAATCCCGCTTAACCCTAT<br>GATGATTGCCGCGTCTTCTATATGTACCGTATTCGGAATAACGGCTTCCGTAGTATT<br>GGATCCTCTTCATGTCTGGCTGTACTGCTTTATTTCGAGTATATTCTTTATATTTGA<br>AATGGTCGTAGCGTTTGCTATATTTGCTATCACGATCCACGATTTTCAGACGATAG<br>GATCACCAATGTCGCTTAAAGTGGTTGAAAGACTCAAACTTATGCGAAgtaagtttaa<br>acatgattttactaactaactaatctgatttaaattttcagTTGTATTCTACGTTTCCTGGATGGCC<br>TATCCTATTCTCTGGTCATTCTCTTCCACGGGTGCTTGTATTATGTCGGAGAACACT<br>TCTTCGGTTTTATATCTTTTGGGCGATGCTCTCTGTAAAAACACGTATGGTATTCT<br>GCTTTGGGCTACTACTTGGGGCCTCCTTAACGGCAAATGGGACCGAGACTATGTT<br>AAGGGCCGAAACGTGGATGGAGCCGCCGCCGTGAGCAAGGGAGAGGAGCTGTT<br>CACCGGAGTGGTGCCCATCCTGGTGGAGCTGGATGGCGACGTGAACGGCCACAA<br>GTTCTCGGTGAGCGGAGAGGGAGAGGGCGACGCCACCTACGGCAAGCTGACCC<br>TGAAGTTCATCTGCACCACCGGCAAGCTGCCCGTGCCGTGGCCAACCCTGGTGAC<br>CACCTTCGGCTACGGCCTGCAGTGCTTCGCCCGCTACCCAGATCACATGAAGCAG<br>CACGACTTCTTCAAGTCGGCCATGCCGGAGGGATACGTGCAGGAGCGCACCATC<br>TTCTTCAAGGATGACGGCAACTACAAGACCCGCGCCGAGGTGAAGTTCGAGGGC<br>GATACCCTGGTGAACCGCATCGAGCTGAAGGGCATCGATTTCAAGGAGGACGGC<br>AATATCCTGGGCCACAAGCTGGAGTACAACTACAATAGCCACAACGTGTACATCAT<br>GGCCGACAAGCAGAAGAACGGCATCAAGGTTAATTTCAAGATCCGCCACAATATC<br>GAGGATGGCTCCGTGCAGCTGGCCGACCACTACCAGCAGAACACCCCGATTGGC<br>GATGGACCCGTGCTGCTgtaagtttaaacatatatatactaactaaccctgattatttaaattttcag<br>GCCAGACAATCACTACCTGAGCTACCAGTCCGCCCTGTCGAAGGACCCCAACGAG<br>AAGCGCGACCACATGGTGCTGCTGGAGTTTGTGACCGCCGCCGGAATTACCCTG<br>GGAATGGACGAGCTGTATAAGttctgctacgagaacgaggtgTAAGAATTCcaactgagcgc<br>cggtcgcta | RAI77_gBlock_ACR1 |
| acccttgGCTAGCgtcgacGGTACCggtagaaaaaATGCCATTCTACGACAGTAGACCGC<br>CGGAAGGTTGGCCAAAGGGTTCCATCAATGATATGGACTACCCGCTCCTCGGTTC<br>CATCTGCGCCGTCTGTTGCGTTTTCGTGGCGGGAAGTGGTATATGGATGCTGTAC<br>CGTTTAGATCTGGGTATGGGATACTCTTGTAAACCGTACAAATCGGGCCGAGCGC<br>CAGAGGTCAATTCTCTGTCCGGAATTATATGTCTGCTTTGCGGCACGATGTATGCG<br>GCGAAATCATTTGATTTCTTTGACGGCGGAGGAACTCCTTTCTCCCTGAACTGGT<br>ATTGGTATCTGGATTATGTGTTCACTTGTCCGCTGCTGATCTTAGATTTTGCGTTCA<br>CATTGGACCTCCCACATAAGATTAGATACTTCTTCGCTGTTTTTTTGACCCTCTGG<br>TGCGGCGTTGCGGCGTTTGTTACACCGAGTGCATACCGTTTCGCGTACTACGCAT<br>TGGGATGCTGCTGGTTCACCCCATTCGCCCTCTCCCTTATGCGACACGTGAAAGA<br>GCGATATTTAGTATACCCGCCGAAGTGTCAGAGATGGCTCTTCTGGGCATGTGTG<br>ATATTCTTCGGTTTTTGGCCGATGTTTCCGATTTTATTCATATTCAGTTGGTTGGGT<br>ACTGGCCATATATCTCAACAGGCTTTCTACATAATCCACGCATTCTTGGACTTAAC<br>GTGTAAATCGATTTTTGGCATATTGATGACTGTATTTCGTCTCGAGTTAGAGGAGC<br>ACACGGAAGTGCAAGGACTGCCTCTTAATGAACCAGAAACCTTATCGGCTGCTGC<br>GAAGTCTCGTATCACATCAGAGGGTGAATACATACCTCTGGATCAAATTGACATAA | RAI321_gBlock_KCR1 |

| | |
|---|---|
| ACGTTgccgccgccaagagcaggatcaccagcgagggcgagtacatccccctggaccagatcgacatcaacgtgGTGAGCAAGGGAGAGGAGCTGTTCACCGGAGTGGTGCCCATCCTGGTGGAGCTGGATGGCGACGTGAACGGCCACAAGTTCTCGGTGAGCGGAGAGGGAGAGGGCGACGCCACCTACGGCAAGCTgtaagtttaaacagttcggtactaactaaccatacatatttaaattttcagGACCCTGAAGTTCATCTGCACCACCGGCAAGCTGCCCGTGCCGTGGCCAACCCTGGTGACCACCTTCGGCTACGGCCTGCAGTGCTTCGCCCGCTACCCAGATCACATGAAGCAGCACGACTTCTTCAAGTCGGCCATGCCGGAGGGATACGTGCAGGAGCGCACCATCTTCTTCAAGGATGACGGCAACTACAAGACCCGCGCCGAGGTGAAGTTCGAGGGCGATACCCTGGTGAACCGCATCGAGCTGAAGGGCgtaagtttaaacatgattttactaactaactaatctgatttaaattttcagATCGATTTCAAGGAGGACGGCAATATCCTGGGCCACAAGCTGGAGTACAACTACAATAGCCACAACGTGTACATCATGGCCGACAAGCAGAAGAACGGCATCAAGGTTAATTTCAAGATCCGCCACAATATCGAGGATGGCTCCGTGCAGCTGGCCGACCACTACCAGCAGAACACCCCGATTGGCGATGGACCCGTGCTGCTGCCAGACAATCACTACCTGAGCTACCAGTCCGCCCTGTCGAAGGACCCCAACGAGAAGCGCGACCACATGGTGCgtaagtttaaacatatatatactaactaaccctgattatttaaattttcagTGCTGGAGTTTGTGACCGCCGCCGGAATTACCCTGGGAATGGACGAGCTGTATAAGttctgctacgagaacgaggtgTGAGAATTCcaactgagcgccggtcgcta | |
| acccttgGCTAGCgtcgacGGTACCggtagaaaaaATGCCTTTTTACGATTCACGTCCTCCGGAAGGCTGGCCTAAGGGCTCCATTAACGATATGGACTATCCGCTGTTGGGCTCAATTTGCGCTGTATGTTGCGTCTTCGTCGCTGGATCGGGTATCTGGATGCTGTATCGACTTGATTTAGGTATGGGCTATTCCTGTAAGCCGTATAAATCAGGACGAGCACCGGAAGTGAACTCTCTTTCAGGCATAATATGTCTCCTTTGTGGTACTATGTACGCCGCAAAGTCATTCGATTTTTTCGACGGAGGCGGCACTCCGTTTTCGCTCAATgtaagtttaaacagttcggtactaactaaccatacatatttaaattttcagTGGTACTGGTATTTAGACTATGTATTCACTTGTCCACTGTTAATCCTCGATTTCGCATTCACATTGGACCTTCCACACAAGATTCGATACTTTTTTGCTGTTTTCTTAACACTCTGGTGTGGCGTGGCGGCCTTTGTAACTCCATCTGCGTATCGATTCGCTTATTACGCACTCGGATGTTGCTGGTTCACACCTTTTGCGCTCAGTCTGATGCGTCACGTTAAAGAGCGATATTTGGTGTATCCTCCTAAGTGCCAACGATGGTTATTCTGGGCCTGTGTGATCTTTTTCGGCTTTTGGCCTATGTTCCCGATACTCTTTATTTTTTCGTGGCTTGGAACCGGCCACATTTCGCAGCAGGCATTTTACATTATTCACGCATTCCTCGACCTCACTTGTAAATCGATCTTTGGAATACTTATGACTGTGTTTAGATTGGAACTGGAGGAGCATACCGAAGTACAGGGCTTGCCATTGAATGAACCGGAGACGTTATCCACGGGAGGAGGTGGAGGATCGGGTGGTGGAGGTTCAGGAGGCGGAGGCAGTGGCTCTACAGGAGGAGGAGGAGGATCAGGAGgtaagtttaaacatgattttactaactaactaatctgatttaaattttcagGAGGAGGATCAGGAGGAGGAGGATCAggaTCAGTGAGCAAGGGAGAGGAGCTGTTCACCGGAGTGGTGCCCATCCTGGTGGAGCTGGATGGCGACGTGAACGGCCACAAGTTCTCGGTGAGCGGAGAGGGAGAGGGCGACGCCACCTACGGCAAGCTGACCCTGAAGTTCATCTGCACCACCGGCAAGCTGCCCGTGCCGTGGCCAACCCTGGTGACCACCTTCGGCTACGGCCTGCAGTGCTTCGCCCGCTACCCAGATCACATGAAGCAGCACGACTTCTTCAAGTCGGCCATGCCGGAGGGATACGTGCAGGAGCGCACCATCTTCTTCAAGGATGACGGCAACTACAAGACCCGCGCCGAGGTGAAGTTCGAGGGCGATACCCTGGTGAACCGCATCGAGCTGAAGGGCATCGATTTCAAGGAGGACGGCAATATCCTGGGCCACAAGCTGGAGTACAACTACAATAGCCACAACGTGTACATCATGGCCGACAAGCAGAAGAACGGCATCAAGGTTAATTTCAAGATCCGCCACAATATCGAGGATGGCTCCGTGCAGCTGGCCGACCACTACCAGCAGAACACCCCGATTGGCGATGGACCCGTGCTGCTgtaagtttaaacatatatatactaactaaccctgattatttaaattttcagGCCAGACAATCACTA | RAI322_gBlock_KCR1(GS) |

| | |
|---|---|
| CCTGAGCTACCAGTCCGCCCTGTCGAAGGACCCCAACGAGAAGCGCGACCACAT<br>GGTGCTGCTGGAGTTTGTGACCGCCGCCGGAATTACCCTGGGAATGGACGAGCT<br>GTATAAGttctgctacgagaacgaggtgGAATTCcaactgagcgccggtcgcta | |
| acccttgGCTAGCgtcgacGGTACCggtagaaaaaATGCCATTCTACGACAGTAGACCGC<br>CGGAAGGTTGGCCACGAGGCTCCGTGAACGACATGGACTATCCGCTTCTGGGCT<br>CAATCTGTGCGATTTCTGCGATAGCTATTGCGGGATCGGGCATATGGATGTTATAT<br>CGATTAGACCTGGGAATGGGCTATTCTTGCAAACCGTATAAATCAGGTCGTGCTC<br>CTGAAGTAAACTCTATTTCCGGAATAGTGTGCCTGCTCTGCGGCACAATGTACGC<br>AGCCAAATCGTTCGATTTTTTCGACGGTGGTGGAACTCCATTTTCACTTAACTGG<br>TACTGGTACTTAGATTATGTGTTCACCTGTCCTCTGTTGATAGTTGATTTCGCGTTC<br>ACCCTGGACATTCCACAAAAGCTGAGATACTCCATTGCCGTATTCGTCGCTCTGTG<br>GTGTGCTGTAGCGGCGTTCGCTACACCATCGGCTTTTCGATTCGCGTATTATGCGC<br>TCGGTTGCTGTTGGTTCATCCCTCTGTCTCTGTTCCTTATACGAGACGTAAAAAG<br>CGTTACCAGGTTTATCCGCCAAAGTGCCAGCGTCTGCTGTTTTGGGCCTGTGTTG<br>TCTTCTTTGGATTCTGGCCTTTGTTTCCATTGCTCTTTATCTTTTCGTGGCAGGGT<br>TCTGGACACATCTCGCGTCAAGCGTATTACATCATCCATGCTTTCCTTGACTTAGT<br>ATGCAAGTCTATATTTGGATTTTTGATGACTTTCTTCCGATTAGAATTAGAGGAGC<br>ACACAGAAGTACAGGGTCTCCCACTGAAGGAACCGAAAGTCATGGACGCTGCGG<br>CTAAATCCCGAATAACGTCTGAAGGAGAGTATATCCCATTGGATCAAATTGACATC<br>AACGTAgccgccgccaagagcaggatcaccagcgagggcgagtacatccccctggaccagatcgaca<br>tcaacgtgGTGAGCAAGGGAGAGGAGCTGTTCACCGGAGTGGTGCCCATCCTGGTG<br>GAGCTGGATGGCGACGTGAACGGCCACAAGTTCTCGGTGAGCGGAGAGGGAGA<br>GGGCGACGCCACCTACGGCAAGCTgtaagtttaaacagttcggtactaactaaccatacatattt<br>aaattttcagGACCCTGAAGTTCATCTGCACCACCGGCAAGCTGCCCGTGCCGTGGC<br>CAACCCTGGTGACCACCTTCGGCTACGGCCTGCAGTGCTTCGCCCGCTACCCAGA<br>TCACATGAAGCAGCACGACTTCTTCAAGTCGGCCATGCCGGAGGGATACGTGCA<br>GGAGCGCACCATCTTCTTCAAGGATGACGGCAACTACAAGACCCGCGCCGAGGT<br>GAAGTTCGAGGGCGATACCCTGGTGAACCGCATCGAGCTGAAGGGCgtaagtttaaa<br>catgattttactaactaactaatctgatttaaattttcagATCGATTTCAAGGAGGACGGCAATAT<br>CCTGGGCCACAAGCTGGAGTACAACTACAATAGCCACAACGTGTACATCATGGCC<br>GACAAGCAGAAGAACGGCATCAAGGTTAATTTCAAGATCCGCCACAATATCGAG<br>GATGGCTCCGTGCAGCTGGCCGACCACTACCAGCAGAACACCCCGATTGGCGAT<br>GGACCCGTGCTGCTGCCAGACAATCACTACCTGAGCTACCAGTCCGCCCTGTCGA<br>AGGACCCCAACGAGAAGCGCGACCACATGGTGCgtaagtttaaacatatatatactaacta<br>accctgattatttaaattttcagTGCTGGAGTTTGTGACCGCCGCCGGAATTACCCTGGGA<br>ATGGACGAGCTGTATAAGttctgctacgagaacgaggtgTGAGAATTCcaactgagcgccggt<br>cgcta | RAI323_gBlo<br>ck_KCR2 |
| ttttcaggaggacccttgGCTAGCgtcgacGGTACCggtagaaaaa | RAI215_ACR<br>1-f |
| ctcagttgGAATTCTTAcacctcgt | RAI216_ACR<br>-1-YFP-r |
| ttttcaggaggacccttgGCTAGCgtcgacGGTACCggtagaaaaaATGCCATTCTACGA | RAI217_KCR<br>-1-f |
| ctcagttgGAATTCTCAcacctcgttctcgtagcagaaCTTATAC | RAI218_KCR<br>-1-YFP-r |

| | |
|---|---|
| GCTAGCgtcgacGGTACCggtagaaaaaATGCCTTTTTACGATTC | RAI219_KCR-1-GS-f |
| ctcagttgGAATTCcacctcgttct | RAI220_KCR-1-GS-YFP-r |
| actgactgGGCCGGCCTTCCTTCAGAAGACGTGCTTTCCTTTTTC | RAI229_snt-1p-FseI-F |
| cctctagaGGCGCGCCGGTGACTGAAAGTTTGATTGATAAATGAA | RAI230_snt-1p-AscI-R |

**Supplementary videos SV1-SV9. Fly behavior before, during, and after light actuation of opsin-bearing flies.**

All videos show *Drosophila* activity before, during and after opsin actuation in the Trumelan activity-monitoring assay (see Methods in the main manuscript). The initial 58 s of each video show flies being exposed to infrared illumination only, followed by 60s of exposure to green light (λ 530 nm) illumination and a second epoch of 60s infrared illumination. In videos SV1-SV3 the flies were maintained on food supplemented with 1mM ATR (see methods in the main manuscript) and a green light illumination intensity of 23.7 μW/mm$^2$ was used. Flies in videos SV4-SV6 were maintained on 2 mM ATR food and the intensity of 44.6 μW/mm$^2$ was used to actuate the opsins. The respective opsins were expressed in motor neurons with *OK371-Gal4*. Upon green light exposure (at the 58s elapsed time point in each video) all ACR1 flies (SV1 and SV4), KCR1-ET flies (SV2 and SV5) and KCR1-GS flies immediately fell on their back and remained in this position until the green light was switched off (1min 58s elapsed time point in each video). At the end of green light illumination (1min 59s elapsed time point in each video) all flies regained their upright posture and displayed unimpaired locomotor activity. Throughout the opsin actuation epoch sporadic limb movement was observed in KCR-expressing flies that was absent in ACR1 flies. Activity was captured at 10 FPS. **SV7-9:** The videos show *C. elegans* activity before, during and after opsin actuation in an open field arena (see methods in the main manuscript). In the initial 10s of each video the worms were exposed to infrared illumination, followed by 10s of green (λ 530 nm, 75 μW/mm$^2$, SV7-SV8) or blue (λ 460 nm, 65 μW/mm$^2$, SV9) light and a second 40s epoch of infrared only illumination. All opsins were expressed pan-neuronally by using the *snt1-p* driver. Prior to the experiment the worms were maintained on media that contained 2mM ATR (see methods in the main manuscript). ACR1 (SV7), KCR1-ET (SV8) and KCR2-ET (SV-9) worms stopped crawling and remained stationary during the opsin actuation period and resumed their crawling behaviour shortly afterwards. Activity was captured at 30FPS.
